# Improved read/write cost tradeoff in DNA-based data storage using LDPC codes

**DOI:** 10.1101/770032

**Authors:** Shubham Chandak, Kedar Tatwawadi, Billy Lau, Jay Mardia, Matthew Kubit, Joachim Neu, Peter Griffin, Mary Wootters, Tsachy Weissman, Hanlee Ji

## Abstract

With the amount of data being stored increasing rapidly, there is significant interest in exploring alternative storage technologies. In this context, DNA-based storage systems can offer significantly higher storage densities (petabytes/gram) and durability (thousands of years) than current technologies. Specifically, DNA has been found to be stable over extended periods of time which has been demonstrated in the analysis of organisms long since extinct. Recent advances in DNA sequencing and synthesis pipelines have made DNA-based storage a promising candidate for the storage technology of the future.

Recently, there have been multiple efforts in this direction, focusing on aspects such as error correction for synthesis/sequencing errors and erasure correction for handling missing sequences. The typical approach is to use separate codes for handling errors and erasures, but there is limited understanding of the efficiency of this framework. Furthermore, the existing techniques use short block-length codes and heavily rely on read consensus, both of which are known to be suboptimal in coding theory.

In this work, we study the tradeoff between the writing and reading costs involved in DNA-based storage and propose a practical scheme to achieve an improved tradeoff between these quantities. Our scheme breaks with the traditional separation framework and instead uses a single large block-length LDPC code for both erasure and error correction. We also introduce novel techniques to handle insertion and deletion errors introduced by the synthesis process. For a range of writing costs, the proposed scheme achieves 30-40% lower reading costs than state-of-the-art techniques on experimental data obtained using array synthesis and Illumina sequencing.

The code, data, and Supplementary Material is available at https://github.com/shubhamchandak94/LDPC_DNA_storage.

## 1 Introduction

In recent years, the amount of data being generated and stored is increasing at rapid rates. As a result of the impending data crisis where generation exceeds reasonable storage capacity, there has been significant interest in exploring alternatives to solid state disks and magnetic tapes as data storage media. Interestingly, the cost of DNA sequencing has been decreasing exponentially in the past ten years. DNA is a robust method of storing information as demonstrated by every living organism and offers exceptionally high storage densities (100s of Petabytes per gram [1]) and long-term durability (1000s of years [2]). The longevity of DNA-based storage makes it ideal as an archival medium to store the knowledge gained by humanity over the millennia. Along with high density data storage, DNA-based storage systems allow efficient duplication of data and random access using PCR-based techniques [3][4].

DNA-based storage involves encoding data into DNA sequences, synthesizing these sequences and later reading them back using sequencing technologies. Since both the DNA synthesis and sequencing processes are inherently error-prone, error correction coding based on the characteristics of this noise is critical for reliable decoding of data. There has been a series of recent works on DNA-based storage such as [1], [4], [5], [6], [7] and [8] which focus on error correction and random-access retrieval, among other aspects.

Figure 1 shows a schematic of a typical DNA-based storage system. Binary data is encoded in the form of short DNA sequences (oligonucleotides) of length around 150 bases where each nucleotide belongs to the set {A, C, G, T}. We assume that the binary data is a uniformly random stream, which can be obtained by lossless compression and encryption of the actual data. The currently standard synthesis process generates millions of copies of each oligonucleotide [9], possibly with errors – substitutions, insertions and deletions. The first step of the reading process involves amplification of the synthesized DNA sequences using PCR [3]. The amplified product is then “sequenced” by randomly sampling oligonucleotides and reading them, possibly with additional errors (usually substitutions for Illumina sequencing). The sequenced oligonucleotides are usually referred to as “reads”. Ideally, the sampling can be modelled as Poisson random sampling, but in practice there is additional sampling bias introduced in the synthesis as well as in PCR amplification performed before sequencing [10]. Due to skewed sampling or bias introduced through synthesis and molecular amplification, some oligonucleotides might be read multiple times while other oligonucleotides might not be read at all. Furthermore, the ordering of the oligonucleotides is completely lost in the process and the synthesized oligonucleotides form an unordered set of DNA sequences. The encoding and decoding algorithms need to handle all these sources of error to recover the data.

**Figure 1:**
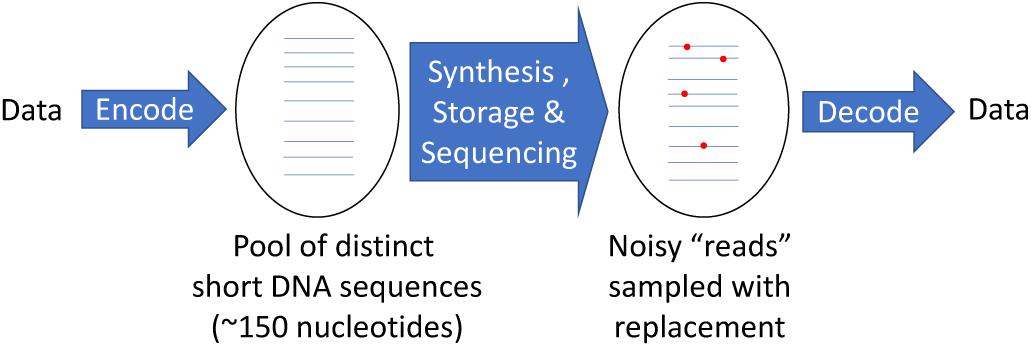
Schematic for a DNA-based storage system. The encoding process converts data to a pool of DNA sequences, typically applying error correction techniques and appending an index to the DNA sequences to store their order. These sequences are then synthesized, stored for some duration and sequenced back to give a pool of noisy reads from the encoded pool. The decoder recovers the data from these noisy reads using error correction techniques.

### Previous work

Over the past years, DNA-based storage has been studied both from practical and theoretical perspectives. In most works, the data is split into short segments at some step in the encoding process due to limits imposed by scalable synthesis techniques. Since the ordering of these segments is lost after synthesis, the typical approach involves appending an index to each oligonucleotide to recover its position in the encoded data stream. Error correction coding is performed to correct errors within each oligonucleotide and to deal with missing oligonucleotides. Two of the state-of-the-art works [1] and [4] use very distinct schemes for the error correction and decoding and are discussed below.

In [1], Fountain codes [11] are used to recover missing oligonucleotides, and Reed Solomon (RS) codes [12] are used to correct errors within each read. Since the RS code is applied on very short sequences, the scheme suffers from limitations of short block length codes [13]. The scheme also ignores reads with insertions and deletions, which usually comprise around 30-50% of the reads [9].

The work in [4] uses large block length RS codes to correct both erasures and errors. However the RS codes operate on symbol-level errors (16-bit symbols in their work) and hence are not optimal for single base errors (i.e. substitutions) that may be introduced by Illumina sequencing. To correct insertion and deletion errors, they cluster the reads by similarity and then apply multiple sequence alignment (MSA) [14] to obtain a consensus sequence. While the consensus allows them to handle some reads with insertions and deletions, it requires multiple reads for a given oligonucleotide and is suboptimal in terms of the sequencing cost. They also attempt to reduce the synthesis error rate by avoiding repeated bases (e.g., AA) using a run-length constraint on the oligonucleotides. This run-length constraint also provides some level of error detection at the oligonucleotide level, but it is likely not the optimal code for this purpose, given that it adds around 25% redundancy to the oligonucleotides. We should note here that their scheme was tested for both Illumina and nanopore sequencing, while this work focuses on Illumina sequencing.

While previous works have proposed several coding schemes, there has been little understanding of the optimal tradeoff between writing cost (bases synthesized/information bit) and reading cost (bases sequenced/information bit). Recent work in [15] and [16] studied the information-theoretic capacity of a DNA-based storage channel, however the work has limited practical applicability due to various unrealistic assumptions and the asymptotic nature of their results.

### Our contributions

In this work, we first analyze the fundamental quantities associated with DNA-based storage systems and understand the associated tradeoffs by theoretical analysis and simulations. Based on this assessment, we propose a practical and efficient scheme to achieve an improved tradeoff between writing cost and reading cost by combining ideas from modern coding theory such as large block length LDPC codes [17] with heuristics to handle insertions and deletions.

Section 2 motivates the proposed approach using a simplified theoretical model. Building upon this, we present the complete encoding and decoding algorithms in Section 3. In Section 4, we present the results for real data (data obtained from synthesis and sequencing experiments) and also discuss the impact of various parameters and synthesis/sequencing non-idealities using this data as well as simulations.

## 2 Theoretical analysis

In this section, we consider a simplified model for DNA-based storage to develop a better understanding of the coding theoretic tradeoffs. While several previous works such as [15], [16], [18] theoretically analyze various aspects of the DNA-based storage problem (such as the information-theoretic capacity in the asymptotic setting and the optimality of various techniques to recover the order of the oligonucleotides), our main focus is to understand the tradeoff between the writing and reading cost associated with DNA-based storage and to motivate the scheme described in Section 3.

### Model and notation

Figure 2 shows the simplified storage model where *nL* message bits are encoded as *nc*_w_ binary sequences of length *L* bits each. We sequence *nc*_r_ reads of length *L* which are passed through a binary symmetric channel with error probability *ϵ*. Here *c*_w_ denotes the average number of encoded bits synthesized per information bit and *c*_r_ denotes the average number of encoded bits read per information bit for successful decoding. Clearly both *c*_w_ and *c*_r_ must be at least 1.

**Figure 2:**
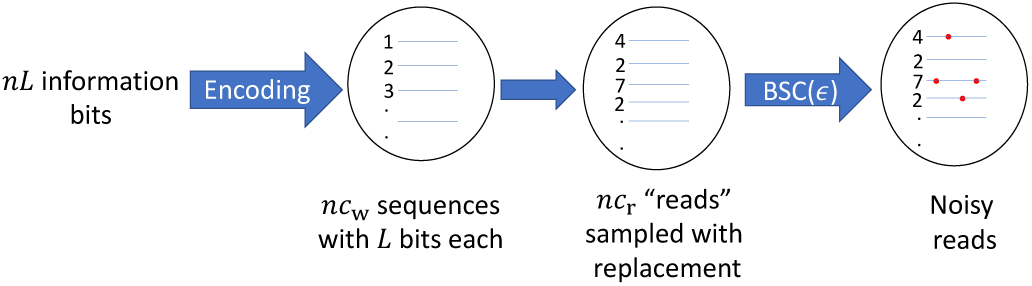
Schematic for simplified storage model. Here *L* denotes the length of synthesized sequences, *c*_w_ denotes the writing cost, *c*_r_ denotes the reading cost and *ϵ* denotes the error rate. The index of each sequence is shown to its left, and the decoder has direct access to the index in the simplified model.

#### Remark

Most previous works in DNA-based storage use coverage (defined as the average number of bits read per synthesized bit = *c*_r_*/c*_w_) as the metric instead of the reading cost *c*_r_. We use *c*_r_ because it captures the actual cost of reading per *information* bit, unlike coverage which measures cost of reading per *encoded* bit and hence coverage comparisons across systems with different *c*_w_ are not meaningful.

For simplicity, we work with bits instead of bases but the results can be extended to an arbitrary alphabet. Furthermore, we assume that the decoder has direct access to the “index” of each read, i.e., the decoder knows the position of the encoded sequence corresponding to any given read (the numbers shown in Figure 2). This simplifies the analysis considerably and can be achieved in a practical system by attaching the index (possibly with error correction) to each sequence (see Section 3). As the theory of channels with insertions and deletions is not adequately understood [19], we ignore insertion and deletion errors in the theoretical analysis for simplicity. In practice, we deal with insertions and deletions by converting them to substitution errors/erasures using MSA and synchronization markers (see Section 3). Finally, we consider an ideal Poisson random sampling model. This model does not capture the synthesis and sequencing bias seen in practice, but while this assumption changes the absolute values obtained in the analysis, it should not affect the tradeoffs and comparisons. As we see below, we can gain significant insights about the real system despite these assumptions.

### Theoretical bounds on read/write cost tradeoff

We first compute the optimal tradeoff between *c*_w_ and *c*_r_ when *ϵ* = 0, i.e., the reads are error-free. In this case, for large enough *n*, we can use the Poisson(*λ*) approximation for the number of times each sequence is observed with *λ* = *c*_r_*/c*_w_. Since the probability of seeing zero copies of a sequence is *e*^*−λ*^, this gives us an erasure channel with capacity 1 *− e*^*−λ*^ [20]. For reliable recovery, we need that the rate *1/c*_w_ be less than the capacity. This gives us

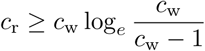

We see that *c*_r_ decreases monotonically with *c*_w_ (Figure 4), implying a tradeoff between the reading and writing costs. This tradeoff can be explained by the fact that decoding *nL* message bits successfully requires at least *n* reads of length *L* with distinct indices. As *c*_w_ increases, random sampling is less likely to produce repeated reads and we need fewer reads to obtain *n* reads with distinct indices.

When *ϵ* ≠ 0, we can obtain lower bounds on the capacity by considering a memoryless approximation to the channel where the input is a bit and the output is a tuple (*k*_0_, *k*_1_) (*k*_*b*_ is the number of times that the bit is read as *b*). The transition probability for this channel is given by

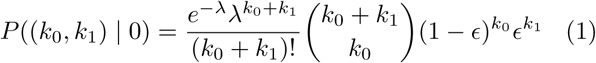

which indicates the probability that a Poisson random variable with parameter *λ* = *c*_r_*/c*_w_ takes value *k*_0_ + *k*_1_ and that we have *k*_1_ errors (for input = 1, swap *k*_0_ and *k*_1_). This is a binary input symmetric channel and hence the capacity achieving distribution is the uniform distribution on the inputs [20]. The capacity and the tradeoff can be numerically computed (see Figure 4 for *ϵ* = 0.5%). For higher *c*_w_ and more powerful error correction, the dependence on consensus decreases and we need fewer copies of each sequence for successful decoding. Note that the bound obtained here need not be tight since the bits are read as entire sequences, not individually. The formulation still provides an asymptotically achievable bound for the simplified model since we can always shuffle the bits at the encoder and the decoder to get a memoryless channel.

### Comparison of coding strategies

We consider two general strategies (shown in Figure 3):

1. **Inner/outer code separation:** Here we first segment the data, apply an erasure-correcting outer code to the segments and then apply an error-correcting/detecting inner code to each segment. During decoding, we first collect the reads corresponding to the same index and take a majority vote at each position. Then we apply the inner code decoding on each segment followed by the outer code decoding to obtain the decoded data. Since *L* is generally small (a few hundred bits), this strategy suffers from the fundamental limits on short block length codes [13] for the inner code. On the other hand, near-optimal erasure-correcting outer codes such as RaptorQ codes [21] are available since *n* can be large.
2. **Single large block code:** Here we apply a single code designed for the channel described in (1) to the data and then segment the encoded data. The decoding just collects the counts (*k*_0_, *k*_1_) at each position using the index of the reads and applies the decoding for the code. For a capacity-achieving code, this strategy will approach the tradeoff curve described earlier as *n* → ∞.

**Figure 3:**
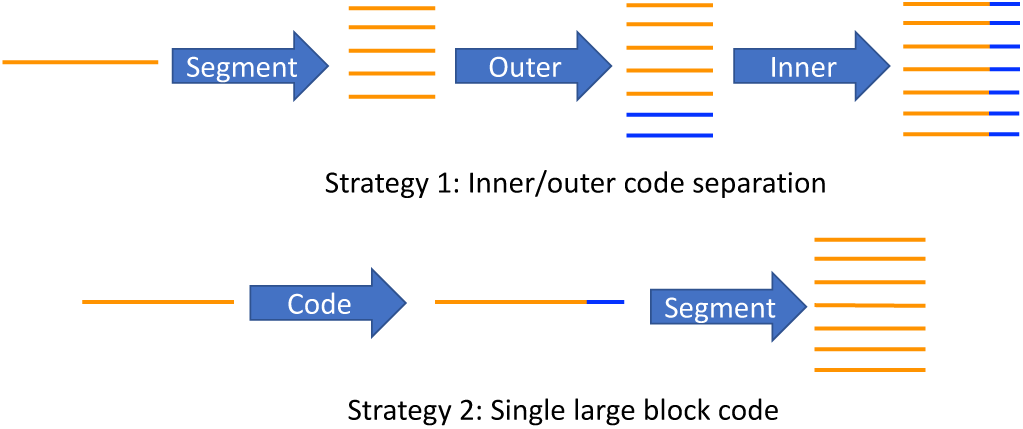
Two general strategies for DNA-based storage. The first strategy uses separate inner and outer codes for correcting erasures of sequences and errors within sequences, while the second strategy just uses a single large block code.

Most previous works on DNA-based storage use the separation strategy, sometimes incorporating elements of the large block code strategy. The work in [11] follows the separation strategy with Fountain codes as the outer codes and RS codes as inner codes. The work in [4] combines the two strategies to some extent by using a run-length constrained code as the inner code (only error detection) and RS code as the outer code that can handle both errors and erasures. However, the run-length constraint in [4] is quite strict (25% added redundancy) and thus the outer code is involved mostly in erasure correction. Also, the RS code in [4] handles errors at a 16-bit block level which is suboptimal for point substitution errors. Finally, there are some works [22], [23] using large block codes such as LDPC codes in the context of DNA-based storage, but they focus on applying LDPC codes separately on each segment and hence follow the separation strategy.

Figure 4 shows some simulations for the two strategies along with the bound for *ϵ* = 0.5% (typical substitution rate for Illumina sequencing). We set the parameters *n* = 1000 and *L* = 256 bits (corresponding to 128 bases). For the separation strategy, we use BCH codes [24] as the inner codes and Raptor codes [21] as the outer codes. Both these codes are among the best known for the parameters in question. For the large block code strategy, we use a LDPC code with a message block of 256,000 bits. We see that the large block code strategy is closer to the bound and outperforms the separation strategy, with close to 2x lower reading cost for low writing costs. More details about these simulations are available in Supplementary Material Section 2.

**Figure 4:**
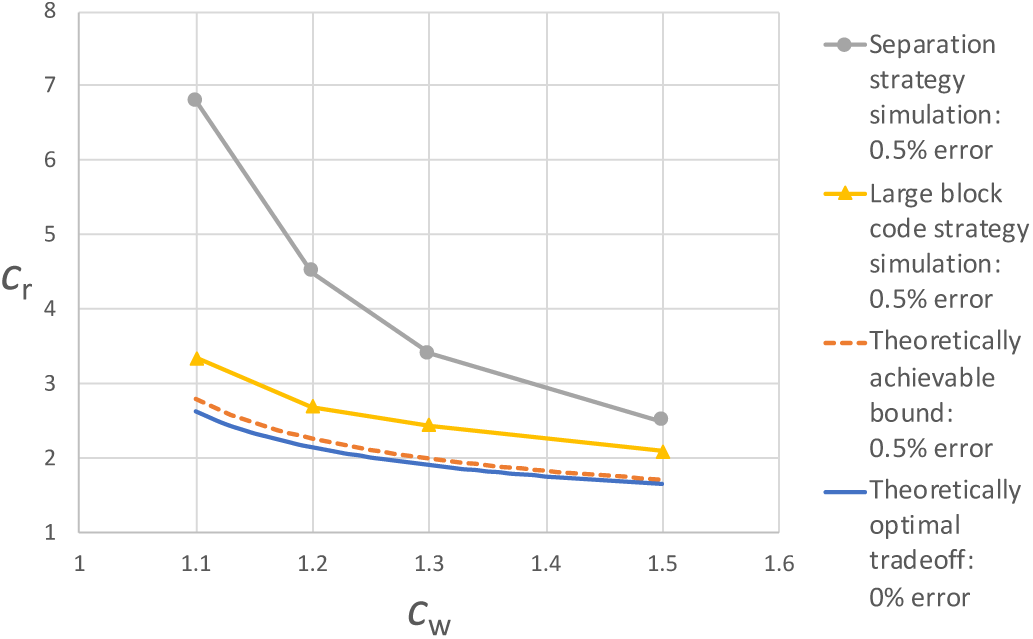
Tradeoff between reading cost *c*_r_ and writing cost *c*_w_ in the simplified model: theoretical bounds for 0% and 0.5% error rates along with simulation results for two strategies at 0.5% error rate: (i) inner/outer code separation strategy (BCH+Raptor codes) and (ii) single large block code strategy (LDPC code).

The analysis and simulations presented here for the simplified model suggest that using a large block length code can provide significant benefits as compared to a inner-outer code separation strategy. In Section 3, we build upon this idea to develop a DNA-based storage framework that can handle the various non-idealities mentioned earlier. The results in Section 4 show that the basic intuition developed here holds true for real experiments.

## 3 Methods

In this section, we present the methods used for real experiments, combining ideas from the analysis in the previous section with additional techniques to handle the errors in the DNA-based storage system. We add an error-protected addressing index to each oligonucleotide in order to determine its position in the encoded data. To resolve the issue of missing oligonucleotides and substitution errors, we use a large block length LDPC code [17]. Since the synthesis process also introduces insertions and deletions, we use synchronization markers in each oligonucleotide. We next describe the encoding and decoding scheme in the proposed framework (shown in Figure 5). The impact of some of these elements on the performance are discussed in Section 4.1.

**Figure 5:**
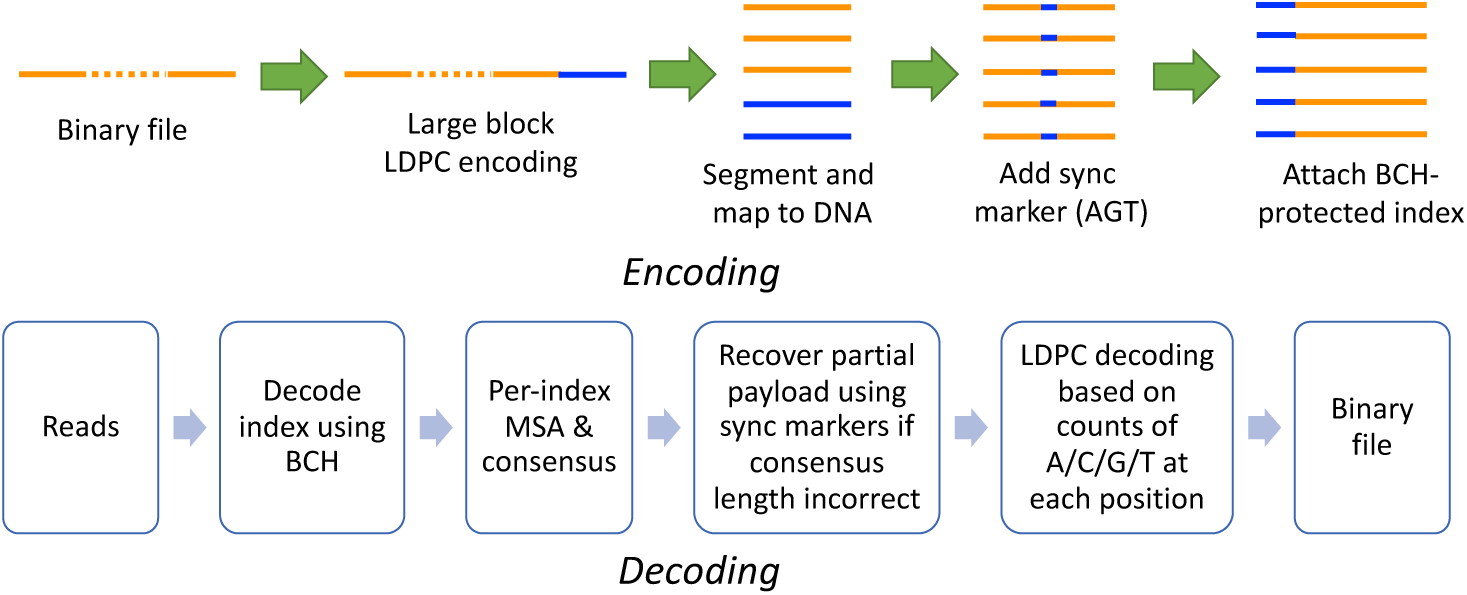
Schematic for proposed scheme.

### 3.1 Encoding

- **LDPC encoding:** Our proposed scheme first encodes the binary data using a large block length LDPC code (in blocks of 256K bits). We use regular LDPC codes from https://github.com/radfordneal/LDPC-codes with a parameter that determines the percentage redundancy. Large block length regular LDPC codes are known to achieve near-optimal performance for a range of channels, including substitution errors and erasures, especially at high rates. The parameters associated with the LDPC codes and the process for encoding and decoding are described in Supplementary Material Section 2.1.
- **Binary to DNA mapping:** These binary blocks are then segmented and mapped to the alphabet {A, C, G, T}. We use the default mapping from 2 bits to a base (00-A, 01-C, 10-G, 11-T). The size of these segments is determined by the capability of the synthesis technique (typically 150-200 bases). Other works ([1], [4]) explored mappings that restrict homopolymers (e.g., AA) to reduce the synthesis error rate at the cost of higher redundancy. Instead, we randomize the binary data itself by compressing and encrypting it before the encoding. Even though this doesn’t completely eliminate homopolymer sequences, we observed that this gives good synthesis quality without increasing the writing cost.
- **Synchronization marker:** We then add a synchronization marker (we use the sequence “AGT”) to the center of each oligonucleotide. These markers are used to recover parts of the oligonucleotide in cases of insertions and deletions as described in Section 3.2.
- **Addressing index:** Finally we add an addressing index to each oligonucleotide. The index is protected with a BCH code [24] (using the code provided at https://github.com/jkent/python-bchlib) that can correct up to 2 bit errors (for the default setting). In our case, the BCH code requires 6*k* bits of redundancy to correct *k* bit errors. It is known that sequences with long homopolymer sequences (e.g., GGGGGG) and abnormal GC content have issues with synthesis/sequencing [1]. To avoid having multiple problematic sequences in the same LDPC block, we also apply a pseudorandom permutation to each index before encoding.

Figure 6 shows the schematic of the encoded oligonucleotides and typical sizes for each component.

**Figure 6:**
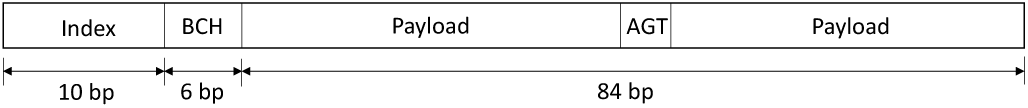
Schematic for the encoded oligonucleotides showing the index with BCH error protection, the payload, and the synchronization marker (AGT) along with their typical sizes.

### 3.2 Decoding

- **Index decoding:** During the decoding process, we first attempt to decode the index of each read using the BCH code. If the decoding fails, we attempt to recover from a single insertion or deletion error in the index. The recovery succeeds when a unique single insertion or deletion error leads to a noise-less BCH codeword. This additional step typically recovers another 5-10% indices (see Section 4.1.4).
- **MSA:** Next, we use Kalign 2 [14] to perform multiple sequence alignment (MSA) for the subset of reads corresponding to each index. As opposed to [4] where the reads are clustered based on similarity, we cluster reads based on their index which is computationally more efficient. If the consensus sequence does not have the correct length, we attempt to recover part of the oligonucleotide using the synchronization marker. For example, if the synchronization marker is shifted left by 1 base, then we retain only the right half of the sequence and consider the left half as an erasure for the next step in the decoding. This allows us to work with very low coverages where we obtain a single erroneous read for a significant fraction of the oligonucleotides.
- **LDPC decoding:** The MSA step provides the counts of 0’s and 1’s for each position in the oligonucleotide. Rather than using just the consensus sequence, we utilize the counts for LDPC decoding by converting the counts into log-likelihood ratios (LLR) using an appropriate probabilistic error model. Denote the substitution error rate as *ϵ* and the counts of 0’s and 1’s at a particular position as *k*_0_ and *k*_1_, respectively. Using this notation, we can compute the log-likelihood ratio as follows (based on (1)):

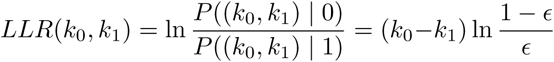 The substitution error rate *ϵ* should be set according to the error rate after consensus, and was set to 4% for our experiments. This is higher than typical substitution error rates from synthesis/sequencing to handle cases like one insertion and one deletion in a single read leading to multiple erroneous bits.

## 4 Experimental results and discussion

To test the performance of the proposed algorithm, we performed nine experiments with different parameters over a period of five months. The synthesis was done by CustomArray (http://www.customarrayinc.com/). The first two experiments were done on separate 12K oligonucleotide pools, while the remaining seven experiments were done in a single 90K pool. The 150 length oligonucleotides consisted of primers of length 25 on either end, which are used for PCR amplification and can also be used for random access. These were then sequenced with Illumina iSeq technology (considering only the first read in a read pair). The detailed experimental procedure is described in Supplementary Material Section 4. Before running the decoder, we used FLEXBAR [25] to remove the primers and detect reverse complemented reads. We encoded a variety of files in these experiments, including random files, an image file (Figure 7) and texts such as the UN declaration of human rights, Darwin’s Origin of Species and Feynman’s speech “There’s Plenty of Room at the Bottom” [26]. To avoid long repeats/homopolymers in the oligonucleotides, the files were randomized using compression and encryption. Details about the experimental parameters and the encoded files are available in Supplementary Material Section 3. The code is available at https://github.com/shubhamchandak94/LDPC_DNA_storage.

**Figure 7:**
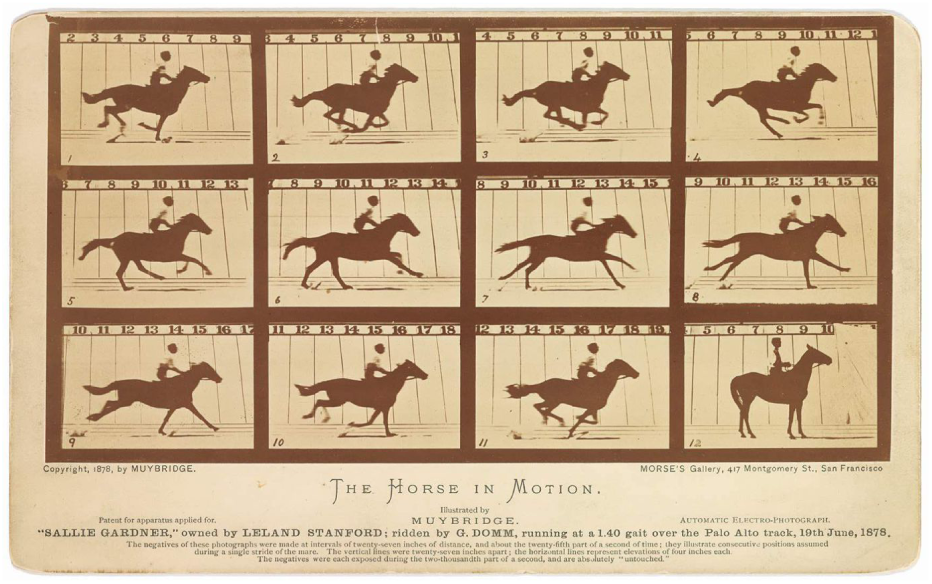
The Horse in Motion image (https://www.loc.gov/pictures/item/97502309/) encoded in the DNA oligonucleotides.

Table 1 shows the results for selected experiments. The remaining experiments explored the impact of secondary parameters such as the BCH code and synchronization marker, and are described in the Supplementary Material. For each experiment, we randomly subsample the reads and find the minimum number of reads for which decoding succeeds in 20 out of 20 trials (multiple trials conducted to ensure robustness – also see Section 4.1.5). The last two columns show the writing cost (bases synthesized/information bit) and reading cost (bases sequenced/information bit). Since a base can represent at most 2 bits, both of these quantities are lower bounded by 0.5. As discussed in Section 2, reading cost represents the actual cost of sequencing more accurately than coverage (bases sequenced/bases synthesized), especially when comparing across different writing costs.

**Table 1:** Results for selected experiments. The first two experiments were synthesized in separate 12K pools while the remaining three were synthesized as part of a larger 90K pool, leading to differences in the coverage variance. The error rates, reading and writing costs are measured excluding the primers. The normalized coverage variance is computed by first subsampling to mean 5x coverage and then normalizing the coverage variance by the variance for ideal Poisson sampling. For ease of comparing results across experiments, only aligned reads were used here (see Section 4.1.2 for details).

| Exp. No. | LDPC redundancy | File size | No. of oligonucleotides | Average error rates |  |  | Normalized coverage variance | Writing cost (bases/bit) | Reading cost (bases/bit) |
| --- | --- | --- | --- | --- | --- | --- | --- | --- | --- |
|  |  |  |  | Substitution | Deletion | Insertion |  |  |  |
| 1 | 50% | 160 KB | 11,710 | 0.39% | 0.86% | 0.05% | 1.97 | 0.91 | 2.73 |
| 2 | 10% | 224 KB | 12,026 | 0.36% | 0.53% | 0.05% | 1.57 | 0.67 | 3.82 |
| 3 | 50% | 192 KB | 13,716 | 0.35% | 0.88% | 0.05% | 3.36 | 0.89 | 3.45 |
| 4 | 30% | 192 KB | 11,892 | 0.47% | 0.85% | 0.05% | 3.53 | 0.78 | 4.46 |
| 5 | 10% | 192 KB | 10,062 | 0.44% | 0.80% | 0.05% | 3.19 | 0.66 | 8.11 |

Table 1 also reports the error rates and the normalized coverage variance for the experiments (see Section 4.1.1 for details). The error rates vary slightly across the experiments, with deletions and substitutions being the most common forms of errors. The normalized coverage variance represents the deviation from Poisson random sampling, with 1 being ideal and higher values representing more variation in the coverage of oligonucleotides. The coverage variance is usually due to a combination of synthesis bias and PCR bias, and can depend on the specifics of the synthesis process [10]. The first two experiments were synthesized separately from the rest and have lower coverage variance. We also note that the first two experiments were conducted before the conception of the synchronization marker idea and thus have slightly different parameters, but we still include the results since they provide valuable insights into the impact of coverage variance and error rates on the performance of the scheme.

From Table 1, we see that as we reduce the LDPC redundancy, the writing cost decreases but the reading cost increases. This is supported by the theoretical analysis done in Section 2. Based on the results, we observe that the higher redundancy (50%) LDPC code is much more resilient to higher coverage variance and error rates as compared to the lower redundancy (10%) LDPC code. This is expected because the 10% LDPC code can correct fewer errors and erasures and hence is impacted more severely if some fraction of oligonucleotides are missing or erroneous.

Table 2 shows the results for two previous works which focused on error correction and reduced writing/reading costs. We note that a direct comparison of our work with the previous works is difficult as the works use different oligonucleotide synthesis providers, encode different amounts of data and use different oligonucleotide lengths. In particular the deletion rate reported in [4] is 0.2% which is significantly lower than the rates in our experiments (around 0.8%). However, experimental results along with the theoretical analysis do suggest that the proposed scheme offers a better tradeoff between the writing and reading costs. In particular, comparing the first two rows of Table 1 with the results in Table 2, we observe that proposed scheme requires 40% lower reading cost at comparable writing costs (also see Figure 8). While the proposed scheme has been tested with smaller file sizes on real datasets, the technique itself is scalable to larger datasets. We discuss the scalability of the proposed technique in Section 4.1.7.

**Table 2:** Results for selected previous works.

| Previous work | File size | No. of oligonucleotides | Writing cost (bases/bit) | Reading cost (bases/bit) |
| --- | --- | --- | --- | --- |
| RS+RLL [4] | 200.2 MB | 13.4 M | 0.91 | 4.55 |
| Fountain+RS [1] | 2.11 MB | 72,000 | 0.65 | 6.80 |

**Figure 8:**
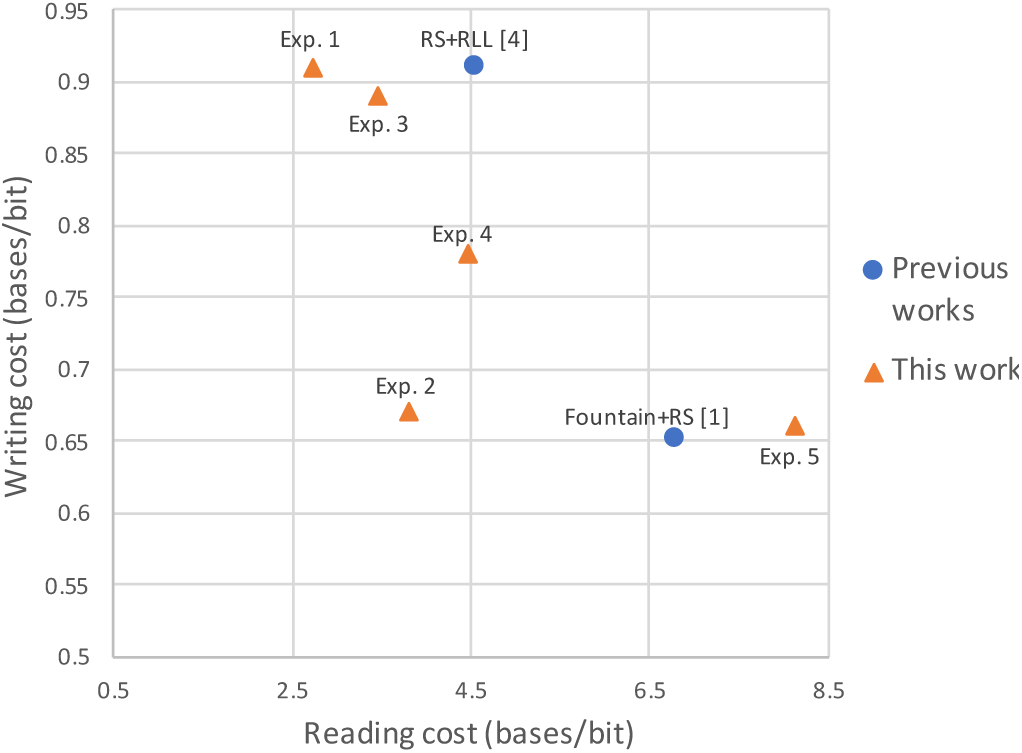
Comparison of results from Tables 1 and 2. The results for this work show variation across experiments due to differences in coverage variance and error rates. In most cases, the proposed approach achieves a better read/write cost trade-off than the previous works in spite of higher deletion rates.

### 4.1 Discussion

In this section, we discuss the error characteristics of the data and the impact of various parameters on the performance of the approach. We also discuss its scalability and perform simulations for stress testing the code with higher error rates. Additional details are available in the Supplementary Material.

#### 4.1.1 Error and coverage profile

Figure 9 shows the substitution, insertion and deletion error rates per position of the 150 length read for experiment #3. These were computed by aligning the reads to the original sequences and include both the synthesis and sequencing errors. The error rate in the primers (first 25 and last 25 bases) is lower because the primers are synthesized separately for performing PCR amplification of the synthesized oligonucleotides. The variation in the deletion rate around positions 60 and 90 are most likely due to the impact of a common synchronization markers across oligonucleotides on the synthesis process (two symmetric peaks due to reverse complementation). The other experiments had similar error profile, although the overall error rate varied a little. On average, most of the experiments had total error rate around 1.3% (substitution: 0.4%, deletion: 0.85%, insertion: 0.05%). Based on the typical error rates for Illumina sequencing and experiments on paired-end data as in [9], the substitution errors are primarily caused by the sequencing and the insertion and deletion errors are primarily from the synthesis.

**Figure 9:**
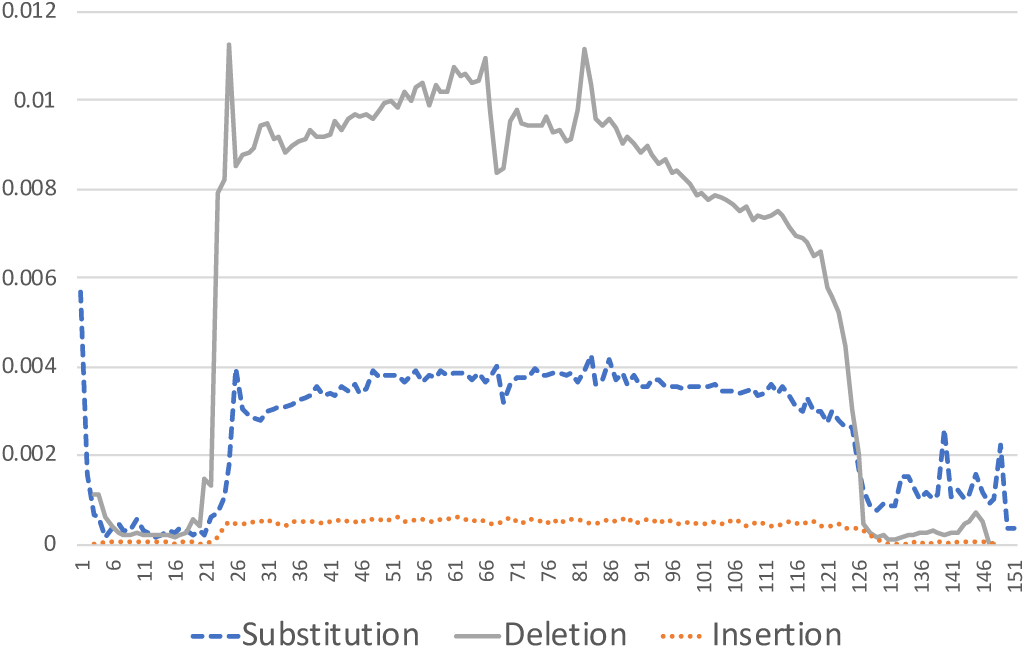
Error rates vs. read position for experiment #3.

Figure 10 shows the histogram of the coverage of the oligonucleotides for experiments #2 and #5 when the mean coverage was set to 5 by subsampling. Here coverage of an oligonucleotide denotes the number of times the oligonucleotide is seen in the subsampled set of reads. The figure also shows the coverage histogram for ideal Poisson sampling at the same mean coverage. We see that the experiments have a larger spread as compared to the ideal sampling and also that experiment #5 had a higher spread than experiment #2 (as seen in the normalized coverage variance from Table 1). In particular, experiment #5 has a significant fraction of oligonucleotides with zero coverage and hence needed a higher reading cost than experiment #2 even though they used the same LDPC redundancy (Table 1). Overall we see that the coverage variance plays a major role in the decoding performance, especially for codes with lower redundancy. We refer the readers to [10] for a detailed analysis of coverage variance in the context of DNA-based storage.

**Figure 10:**
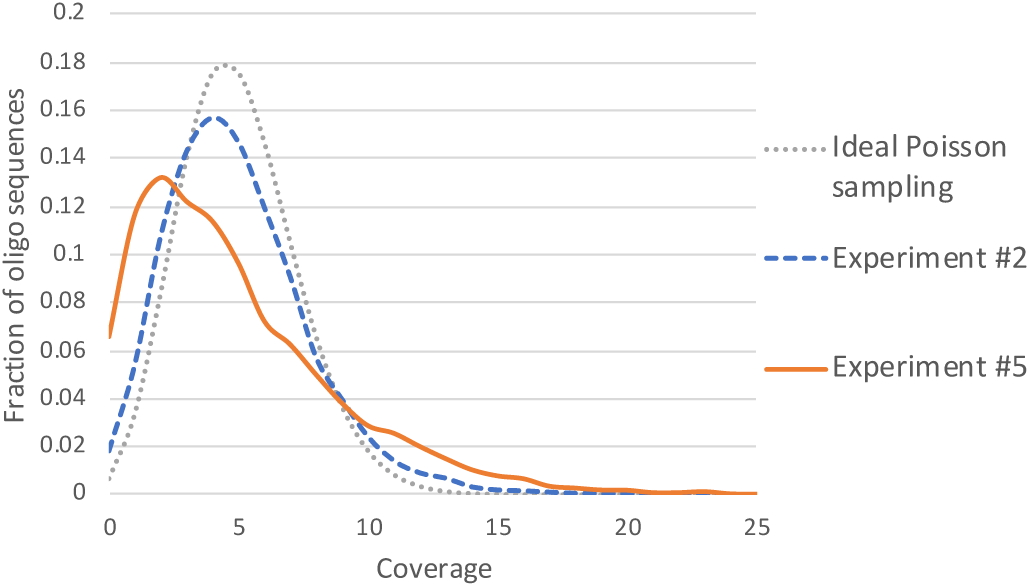
Coverage histogram for ideal Poisson sampling and for experiments #2 and #5 at mean coverage 5.

#### 4.1.2 Unaligned reads in decoding

The sequenced data usually includes some number of reads that do not align properly to the original oligonucleotides. This can be due to synthesis/sequencing errors or due to PhiX spike-in (known PhiX virus DNA added to the sequencing pool for quality control purposes). The decoding algorithm identifies and removes such reads during the primer removal step and the index decoding step. Since the fraction of unaligned varies from experiment to experiment, the results in Table 1 included only the aligned reads for the sake of comparison. If the decoding is performed using all reads, the reading cost increases depending on the percentage of unaligned reads. For example, the first experiment in Table 1 included roughly 13% unaligned reads and the reading cost increases from 2.73 bases/bit to 3.16 bases/bit when these are included in the decoding process.

#### 4.1.3 Impact of BCH code parameter

To understand the impact of the BCH code parameter on the performance of the scheme, we conducted real experiments exploring a range of these parameters. Unfortunately, due to the differences in the coverage variance and error rate across experiments, we were unable to reach any definite conclusion from these experiments (results available in Supplementary Material Section 3). Here we provide some simulation results for these parameters. Simulations are performed with LDPC redundancy 50%, file size 224KB, error rate 1.3% (substitution: 0.4%, deletion: 0.85%, insertion: 0.05%) and with 15% reads being random sequences (to simulate unaligned reads). Table 3 shows the results, where the BCH parameter represents the number of bit errors the BCH code can correct. We use 2 bit error correction because it offers the best balance between writing and reading cost for this error rate.

**Table 3:** Simulation results for various settings of the BCH parameter with realistic error rates. Writing and reading cost are in bases/bit.

| BCH parameter | Writing cost | Reading cost |
| --- | --- | --- |
| 0 | 1.20 | 3.80 |
| 1 | 1.16 | 3.20 |
| 2 | 1.12 | 2.50 |
| 3 | 1.07 | 2.50 |

#### 4.1.4 Impact of insertion and deletion correction heuristics

As discussed in Section 3, we use a couple of heuristics to handle reads with insertions and deletions without relying on consensus. During index decoding, we attempt to correct a single insertion or deletion error with a BCH code which typically can only correct substitution errors. For most files, we observed that this step is able to correct 5-10% additional indexes. Table 4 shows the results with and without this step. We see that this step reduces the reading cost by around 5% for all three LDPC codes without affecting the writing cost.

**Table 4:** Results with and without heuristic for correction of a single insertion or deletion error in the index, for experiments #3-5. Writing cost and reading cost are in bases/bit.

| LDPC redundancy | Writing cost | Reading cost when correcting a single insertion or deletion | Reading cost when correcting no insertions or deletions |
| --- | --- | --- | --- |
| 50% | 0.89 | 3.45 | 3.58 |
| 30% | 0.78 | 4.46 | 4.73 |
| 10% | 0.66 | 8.11 | 8.57 |

We also use a synchronization marker to recover part of the read if the consensus length is incorrect. We conducted real experiments with and without a marker, but as described in Section 4.1.3 the results were not conclusive due to differences in the coverage variance and error rate across experiments. In simulations, we did see that using a marker leads to around 10% improvement in the reading cost while having little impact (2-3%) on the writing cost.

#### 4.1.5 Probability of decoding failure

For most of this work, we have reported the reading cost at which 20 out of 20 sampling trials were successful. This was done for the sake of comparisons and due to computational constraints. In this section, we study the probability of decoding failure as the reading cost is varied. We performed realistic simulations with a single LDPC block (32 KB) and computed the fraction of unsuccessful trials at each reading cost (simulation error model as in Section 4.1.3). Figure 11 shows the plots for three LDPC redundancies. We see that the failure probability falls rapidly from 1 after the reading cost exceeds a threshold, where this threshold is lower for codes with higher redundancy/writing cost. This is consistent with the typical behavior seen in coding theory [17] and suggests that the scheme offers high reliability once the reading cost exceeds the threshold value.

**Figure 11:**
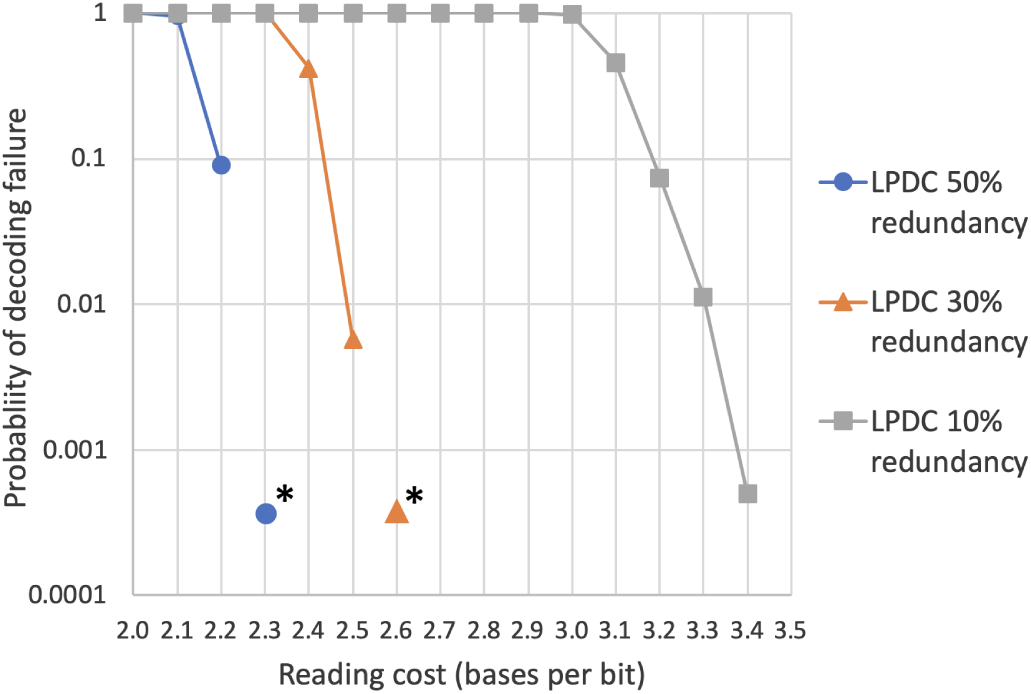
Probability of decoding failure vs. reading cost for realistic simulations for three LDPC redundancies. * denotes that all of the 10,000 trials were successful, suggesting that the failure probability is below the marker shown.

#### 4.1.6 Stress testing

To stress test the framework, we performed simulations at increased error rate of 6% (substitutions, deletions, insertions 2% each) along with 15% reads being random sequences (to simulate unaligned reads). We encoded a 224KB file with 50% redundancy LDPC code and a BCH code capable of correcting 3 errors (writing cost 1.07 bases/bit). Given the high error rate, the *ϵ* parameter for the LDPC decoding (see Section 3) was set to 10% instead of the default value of 4%. The decoding succeeded at a reading cost of 10.5 bases/bit showing that the framework can be used even at relatively high error rates, although it might not be optimal in this setting.

#### 4.1.7 Performance and scalability

The current implementation is written in Python with the libraries for LDPC codes, BCH codes, barcode removal and multiple sequence alignment written in C/C++. All experiments were done on a server with 40-core Intel Xeon processor (2.20GHz) and 256 GB RAM. For the 50% redundancy LDPC code (experiment #3), encoding 224 KB of data takes 1m30s and uses 4.1 GB of RAM. Decoding the corresponding reads takes 56s and uses 190 MB of RAM. Most of the resources are consumed by the single-thread implementation of LDPC encoding and decoding, which can be efficiently implemented in hardware as is done for communication applications. Since the code works in blocks, the time consumption is linear in the file size and the memory consumption of the LDPC coding is constant. Therefore, the proposed scheme is scalable to large files.

## 5 Conclusions and future work

In this work we propose practical and efficient error correcting codes for Illumina sequencing-based DNA-based storage that achieve a better tradeoff between the writing cost and reading cost as compared to previous works. The proposed scheme utilizes ideas from modern coding theory and combines them with heuristics to handle insertion and deletion errors. We believe that the tools, analysis and insights obtained in this project can be useful beyond DNA-based storage to understand the error characteristics of synthesis/sequencing platforms and in developing better bioinformatics algorithms.

Possible future work includes utilizing large block codes optimized for the DNA-based storage channel instead of the regular LDPC codes and using marker codes [27] to handle insertions and deletions in a more systematic way. The improved insertion and deletion correction can extend the applicability of the framework to sequencing platforms such as nanopore sequencing [28] which have higher insertion and deletion error rates. Another interesting direction is to incorporate ideas from [18] and [29] to reduce the inefficiency of index error correction.

## Acknowledgments

We thank Andrea Montanari for helpful discussions.

## References

[1] Y. Erlich and D. Zielinski, “DNA Fountain enables a robust and efficient storage architecture,” Science, vol. 355, no. 6328, pp. 950–954, 2017.

[2] G. R. N. et al., “Robust Chemical Preservation of Digital Information on DNA in Silica with Error-Correcting Codes,” Angewandte Chemie International Edition, vol. 54, no. 8, pp. 2552–2555.

[3] R. K. Saiki et al., “Primer-directed enzymatic amplification of DNA with a thermostable DNA polymerase,” Science, vol. 239, no. 4839, pp. 487–491, 1988.

[4] L. Organick et al., “Random access in large-scale DNA data storage,” Nature biotechnology, vol. 36, no. 3, p. 242, 2018.

[5] G. M. Church et al., “Next-generation digital information storage in DNA,” Science, vol. 337, no. 6102, pp. 1628–1628, 2012.

[6] N. Goldman et al., “Towards practical, high-capacity, low-maintenance information storage in synthesized DNA,” Nature, vol. 494, no. 7435, p. 77, 2013.

[7] M. Blawat et al., “Forward Error Correction for DNA Data Storage,” Procedia Computer Science, vol. 80, pp. 1011–1022, 2016. International Conference on Computational Science 2016, ICCS 2016, 6-8 June 2016, San Diego, California, USA.

[8] S. H. T. Yazdi et al., “A rewritable, random-access DNA-based storage system,” Scientific reports, vol. 5, p. 14138, 2015.

[9] R. Heckel et al., “A characterization of the DNA data storage channel,” CoRR, vol. abs/1803.03322, 2018.

[10] Y.-J. Chen et al., “Quantifying Molecular Bias in DNA Data Storage,” BioRxiv, p. 566554, 2019.

[11] D. J. C. MacKay, “Fountain codes,” IEE Proceedings – Communications, vol. 152, pp. 1062–1068, Dec 2005.

[12] I. Reed and G. Solomon, “Polynomial Codes Over Certain Finite Fields,” Journal of the Society for Industrial and Applied Mathematics, vol. 8, no. 2, pp. 300–304, 1960.

[13] Y. Polyanskiy et al., “Channel coding rate in the finite blocklength regime,” IEEE Transactions on Information Theory, vol. 56, no. 5, pp. 2307–2359, 2010.

[14] O. Frings et al., “Kalign2: high-performance multiple alignment of protein and nucleotide sequences allowing external features,” Nucleic Acids Research, vol. 37, pp. 858–865, 12 2008.

[15] R. Heckel et al., “Fundamental limits of DNA storage systems,” CoRR, vol. abs/1705.04732, 2017.

[16] I. Shomorony and R. Heckel, “Capacity results for the noisy shuffling channel,” CoRR, vol. abs/1902.10832, 2019.

[17] D. J. C. MacKay and R. M. Neal, “Near Shannon limit performance of low density parity check codes,” Electronics Letters, vol. 32, pp. 1645–, Aug 1996.

[18] A. Lenz et al., “Coding over sets for DNA storage,” in 2018 IEEE International Symposium on Information Theory (ISIT), pp. 2411–2415, IEEE, 2018.

[19] M. Mitzenmacher et al., “A survey of results for deletion channels and related synchronization channels,” Probability Surveys, vol. 6, pp. 1–33, 2009.

[20] T. M. Cover and J. A. Thomas, Elements of information theory. John Wiley & Sons, 2012.

[21] A. Shokrollahi, “Raptor codes,” IEEE Transactions on Information Theory, vol. 52, pp. 2551–2567, June 2006.

[22] P. Fei and Z. Wang, “LDPC Codes for Portable DNA Storage,” in 2019 IEEE International Symposium on Information Theory (ISIT), IEEE, 2019.

[23] A. K.-Y. Yim et al., “The essential component in DNA-based information storage system: robust error-tolerating module,” Frontiers in bioengineering and biotechnology, vol. 2, p. 49, 2014.

[24] R. Bose and D. Ray-Chaudhuri, “On a class of error correcting binary group codes,” Information and Control, vol. 3, no. 1, pp. 68 – 79, 1960.

[25] M. Dodt et al., “FLEXBAR—Flexible Barcode and Adapter Processing for Next-Generation Sequencing Platforms,” Biology, vol. 1, no. 3, pp. 895–905, 2012.

[26] R. Feynman, “There’s Plenty of Room at the Bottom, talk given on December 29th 1959,” Sci. Eng, vol. 23, p. 22, 1960.

[27] E. A. Ratzer, “Marker codes for channels with insertions and deletions,” in Annales des télécommunications, vol. 60, pp. 29–44, Springer, 2005.

[28] M. Jain et al., “The Oxford Nanopore MinION: delivery of nanopore sequencing to the genomics community,” Genome biology, vol. 17, no. 1, p. 239, 2016.

[29] A. Lenz et al., “Anchor-Based Correction of Substitutions in Indexed Sets,” arXiv preprint 1901.06840, 2019.

