## Supplementary Material for "Improved read/write cost tradeoff in DNA-based data storage using LDPC codes"

**3 Experimental parameters and results** **3**

**4 Experimental procedure** **4**

### 1 Availability of code and data

The code and instructions for installation and encoding/decoding are available at `https://github.com/shubhamchandak94/LDPC_DNA_storage/` (commit: `42cc1097bad8eee8be776753fffb62ad37e75ff5`). The specific scripts used for running various experiments are mentioned in the text below as relevant. The data (encoded datasets, oligo files sent for synthesis, raw and aligned FASTQ files obtained from sequencing, error statistics and LDPC matrices for decoding) is available at `https://github.com/shubhamchandak94/LDPC_DNA_storage_data/` (commit: `5089d6197ec9354c97b21fea1d1d60a7d87a2b71`).

### 2 Error correcting codes

Error correcting codes [1] are used for transmitting data reliably across noisy channels. The idea is to encode the data (encode message bits to codewords) using some redundancy so that the decoder can recover the original data even if the transmitted signal is corrupted in some way. In this section, we briefly describe the error correcting codes used in our work.

#### 2.1 LDPC codes

Low Density Parity Check (LDPC) codes [2] are a class of linear error correcting codes that can practically achieve reliable communication at rates close to capacity (best achievable rate). LDPC codes are typically decoded using iterative belief propragation decoding which iteratively updates the likelihood ratio of each codeword. The decoding process depends on the channel only through log-likelihood ratios (roughly speaking, the probability of receiving some channel output given the input bit was 0 or 1).

LDPC codes are represented using a bipartite graph with variable nodes (bits in the codeword) and check nodes (representing parity checks), with the check nodes connected to the variable nodes involved in the corresponding parity check equation. This graph is usually constructed randomly according to some distribution on the degrees of the variable and check nodes. Regular LDPC codes have fixed degree variable and check nodes and perform quite well in the high rate regimes for finite block lengths. Irregular LDPC codes can achieve performance closer to capacity but need some fixes to achieve good finite block length performance. Protograph LDPC codes [3] offer the best of both worlds but require some tuning to find the best protograph for the channel at hand. In this work, we use regular LDPC codes for simplicity. A regular LDPC code is represented by two parameters $(d_v, d_c)$ where $d_v$ is the degree of the variable nodes and $d_c$ is the degree of the check nodes. These quantities are related to the redundancy $\alpha$ by the formula

$$\alpha = \frac{d_v}{d_c - d_v}$$

For best performance, $d_v$ is set to 3 and $d_c$ is computed according to $\alpha$. We used the library available at `https://github.com/radfordneal/LDPC-codes` for creating the parity check matrices, generator matrices of the LDPC codes, and also for encoding and decoding. For decoding, we had to add a new mode for our specific channel.

LDPC codes work on blocks of input bits and we set the input block size to 256000 bits to get the benefits of large block lengths while being practical to implement and run. We should note that even though the LDPC block size is quite large, the performance for erasure correction is partly dictated by the fact that the oligos are sampled rather than individual bits, and hence the effective block size for erasures (number of oligos per LDPC block) is smaller. The 256000 bits are encoded into $256000(1 + \alpha)$ bits where $\alpha$ is as defined above. Note that the rate of code (number of input bits/number of encoded bits) is $1/(1 + \alpha)$.

The proposed algorithm requires the LDPC parity check matrix and a vector containing locations of the input bits in the encoded stream (systematic bits) for decoding. The matrices used in our experiment were generated using the script `util/generate_LDPC_matrices.sh` and the matrices are also available at `https://github.com/shubhamchandak94/LDPC_DNA_storage_data/tree/master/matrices`. Encoding also requires access to the code generator matrix which is also generated by the script above.

#### 2.2 BCH code

Our framework uses BCH codes for index error correction. For each $k$, BCH codes can have block length up to $2^k - 1$ bits where $rk$ bits in the block act as the parity bits. Here $r$ is the number of bit errors the BCH code can correct. We used the Python library available at `https://github.com/jkent/python-bchlib` for the implementation. For the index, we used $k = 6$ and hence we needed 6 parity bits for correcting 1 error. Thus, the total index size including the parity bits is limited to 63 in the current implementation. This should be large enough for most practical applications, e.g., it allows 32 bit index (4 billion oligos) with a BCH code capable of correcting 4 bit errors (24 bits parity). By default, we used a BCH code capable of correcting 2 bit errors. Depending on the error rates, this value can be easily changed.

BCH codes are also used as the inner codes in the theoretical analysis section. In that case, we used $k = 8$ to allow oligo length close to 256.

#### 2.3 RaptorQ

In the theoretical analysis section, we use RaptorQ codes [4] (`https://tools.ietf.org/html/rfc6330`) as the outer codes for correcting erasures. RaptorQ codes improve upon Fountain codes [5] and have several desirable properties such as systematic encoding, low complexity and near-optimal erasure correction. For example, $K$ input symbols can be encoded into $N$ encoded symbols (for arbitrary $N > K$) and the decoder is able to decode the input symbols with 99.9999% probability once it receives $K + 2$ unique encoded symbols. In our case each segment can be thought of as a symbol. We used the Python library available at `https://github.com/mk-fg/python-libraptorq` for the implementation.

#### 2.4 Theoretical analysis - simulations

In Figure 4 in the main text, we plot the tradeoff between writing and reading costs obtained from the theoretical analysis along with the performance of two distinct strategies (inner-outer code separation vs. single large block code). For each strategy, we fixed the writing cost by setting the code parameters and increased the reading cost until we achieved success in 100 out of 100 trials. For the inner-outer code separation strategy, we varied both the Raptor and BCH code parameters and chose the combination that gave the lowest reading cost at a given writing cost.

The code for computing the theoretical bounds is available at `util/analysis/capacity_computation/`. The code for the inner outer separation strategy is available at `util/analysis/Raptor_BCH/`. The code for the single large block code strategy is available at `util/analysis/LDPC`.

### 3 Experimental parameters and results

In this section, we describe the parameters used for the real experiments and the corresponding results. Unless otherwise specified, the reading cost was computed by finding the minimum number of reads for which the decoding succeeded for 20 out of 20 random subsampling trials.

| Exp. no. | LDPC redundancy | BCH parameter | Total index length (bases) | Sync marker | Sync position | Encoded file type | Encoded file size (bytes) | No. of oligos | Writing cost (bases/bit) |
|---|---|---|---|---|---|---|---|---|---|
| 1 | 0.5 | 2 | 18 | None | - | Random | 160,000 | 11,710 | 0.91 |
| 2 | 0.1 | 2 | 18 | None | - | Random | 224,000 | 12,026 | 0.67 |
| 3 | 0.5 | 2 | 13 | AGT | 56 | Image | 191,776 | 13,716 | 0.89 |
| 4 | 0.3 | 2 | 13 | AGT | 56 | Image | 191,776 | 11,892 | 0.78 |
| 5 | 0.1 | 2 | 13 | AGT | 56 | Image | 191,776 | 10,062 | 0.66 |
| 6 | 0.5 | 3 | 16 | AGT | 58 | Image | 191,776 | 14,232 | 0.93 |
| 7 | 0.5 | 1 | 10 | AGT | 55 | Image | 191,776 | 13,248 | 0.86 |
| 8 | 0.5 | 2 | 13 | None | - | Image | 191,776 | 13,248 | 0.86 |
| 9 | 0.1 | 1 | 10 | None | - | Text | 286,432 | 14,094 | 0.62 |

Table 1: Parameters for the experiments. The first two experiments used a larger index space (24 bits) than the remaining experiments (14 bits).

| Exp. no. | Total no. of aligned reads | Average error rates | | | Normalized coverage variance | Min. no. of reads for decoding | Reading cost (bases/bit) |
|---|---|---|---|---|---|---|---|
| | | Substitution | Deletion | Insertion | | | |
| 1 | 3,956,040 | 0.39% | 0.05% | 0.86% | 1.97 | 35,000 | 2.73 |
| 2 | 5,593,049 | 0.36% | 0.05% | 0.53% | 1.57 | 68,500 | 3.82 |
| 3 | 369,409 | 0.35% | 0.05% | 0.88% | 3.36 | 53,000 | 3.45 |
| 4 | 1,451,454 | 0.47% | 0.05% | 0.85% | 3.53 | 68,500 | 4.46 |
| 5 | 1,203,987 | 0.44% | 0.05% | 0.80% | 3.19 | 124,500 | 8.11 |
| 6 | 368,096 | 0.39% | 0.05% | 0.89% | 3.40 | 58,000 | 3.78 |
| 7 | 100,627 | 0.31% | 0.05% | 0.76% | 5.01 | 55,000 | 3.58 |
| 8 | 689,932 | 0.40% | 0.05% | 0.77% | 3.75 | 57,500 | 3.75 |
| 9 | 535,673 | 0.49% | 0.05% | 0.87% | 3.18 | * | * |

Table 2: Sequencing and decoding results for the experiments. The error rates are measured excluding the primers. The normalized coverage variance is computed by first subsampling to mean 5x coverage and then normalizing the coverage variance by the variance for ideal Poisson sampling.
* For experiment 9 with the weakest error correction, decoding did not succeed for 20/20 trials even at very high reading costs (decoding did succeed for some fraction of trials at reading cost above 10 bases/bit.

Table 1 shows the parameters for the nine experiments we performed. Note that the first two experiments were synthesized in separate 12K pools before some aspects of the algorithm were developed and hence have slightly different parameters. Rest of the experiments were synthesized in a single 90K pool and separated later using alignment (they can also be separated using the distinct primers). The code for synthesizing these sequences is available at `util/encode_files.py` and more details about other parameters such as primer sequences is available at `util/params.py`. The oligo sequences and the files encoded in the experiments are available at `https://github.com/shubhamchandak94/LDPC_DNA_storage_data`.

Table 2 shows the error characteristics and results for the experiments. The code for alignment and generation of statistics is available at `util/align_compute_stats.sh`. The code for decoding is available at util/decode_files.py. The code for decoding experiment 1 with the unaligned reads and the code for decoding without index indel correction are available at util/decode_files_unaligned.py and util/decode_files_no_attempt_indel_cor.py, respectively.

Finally, the simulations for BCH parameters and for stress testing the code were performed with util/run_simulations.py. The simulations for computing the probability of decoding failure were performed with util/run_simulations_block_error_rate.py.

### 4 Experimental procedure

Single-stranded oligonucleotide pools were ordered from GenScript (previously CustomArray, before acquisition). Pools were individually amplified using PCR primers specific to the experiment or subpool of interest using 1X Kapa HiFi PCR ReadyMix (Roche Biosystems), 500 nM PCR primers, and 1 microliter of the subpool under the following conditions: 98°C for 45 seconds; 12 cycles of 98°C for 15 seconds, 60°C for 30 seconds, and 72°C for 30 seconds; and a final extension step of 72°C for one minute. Each PCR reaction was purified with Ampure XP beads (Beckman Coulter) using standard protocols.

The amplicons were then subject to Illumina library preparation using the Kapa HyperPrep library preparation kit (Roche Biosystems) under standard protocols, with each library receiving a specific sample barcode for demultiplexing. These libraries were then PCR amplified with 1X Kapa HiFi PCR Ready mix and 1X Illumina's P5 and P7 primers under the following conditions: 98°C for 45 seconds; 5 cycles of 98°C for 15 seconds, 60°C for 30 seconds, and 72°C for 30 seconds; and a final extension step of 72°C for one minute. The samples were quantified by Qubit fluorometry (Thermo Fisher Scientific), and then loaded for sequencing on the Illumina iSeq system.

### References

[1] T. M. Cover and J. A. Thomas, *Elements of information theory.* John Wiley & Sons, 2012.

[2] D. J. C. MacKay and R. M. Neal, "Near Shannon limit performance of low density parity check codes," *Electronics Letters*, vol. 32, pp. 1645–, Aug 1996.

[3] Y. Fang, G. Bi, Y. L. Guan, and F. C. Lau, "A survey on protograph ldpc codes and their applications," *IEEE Communications Surveys & Tutorials*, vol. 17, no. 4, pp. 1989–2016, 2015.

[4] A. Shokrollahi, "Raptor codes," *IEEE Transactions on Information Theory*, vol. 52, pp. 2551–2567, June 2006.

[5] D. J. C. MacKay, "Fountain codes," *IEE Proceedings - Communications*, vol. 152, pp. 1062–1068, Dec 2005.
